# Putative Pseudolysogeny-Dependent Phage Gene Implicated in the Superinfection Resistance of *Cutibacterium acnes*

**DOI:** 10.1101/2020.12.15.422974

**Authors:** Stephanie Wottrich, Stacee Mendonca, Cameron Safarpour, Christine Nguyen, Laura J. Marinelli, Robert L. Modlin, Jordan Moberg Parker

## Abstract

**Background:** Bacteriophage therapy is a promising option to minimize the risk of treatment-associated antibiotic resistance in *Cutibacterium acnes*. A novel phage (Aquarius) was isolated and analyzed to explore characteristics of *C. acnes* phages that may confer lysis-evasion properties.

**Materials and Methods:** Phage superinfection resistance assays were performed with a range of *C. acnes* phages, which were assayed for pseudolysogeny via phage release assays and episome PCR. Bioinformatics and qRT-PCR were used to identify candidate genes related to observed phenotypes.

**Results:** Assay findings indicated that infected *C. acnes* strains were broadly resistant to superinfection and were capable of forming stable pseudolysogens. A conserved Ltp family-like gene contained protein signatures which may be contributing to phage-mediated superinfection resistance in a pseudolysogeny-dependent manner.

**Conclusions:** *C. acnes* bacteria are capable of harboring phage pseudolysogens, and this phenomenon may result in superinfection resistance, necessitating consideration in targeting optimization of *C. acnes* phage-based therapy.

## Introduction

*Cutibacterium acnes* is a gram-positive bacterium of the human epidermal microbiome. It has been documented widely within human microcomedones, regardless of skin microflora variability.^1–3^ Certain strains of *C. acnes* have been implicated as key contributors to acne vulgaris (i.e. acne).^2,3^ Though generally considered a mild infliction, many individuals affected by acne may suffer from psychosocial problems and physical pain at affected sites.^2^ Additionally, *C. acnes* contributes to severe health complications such as post-operative prosthetic hardware contamination, sarcoidosis, spondylodiscitis, prostate pathologies, and Parkinson’s Disease.^4,5^ Extensive time and effort have been dedicated to the study of *C. acnes* and *C. acnes* bacteriophages to better understand and characterize the predominant strains in humans, particularly those correlated with pathogenicity.^1,3^

In conjunction with these efforts, studies have evaluated the efficacy of phage therapy for preventing morbidities that result from *C. acnes* infection.^6,7^ The investigation of parameters such as biofilm formation have led to the discovery of *C. acnes* resistance to phage infection in some cases, presenting an obstacle to phage therapy.^6,8,9^ The leading proposal for this phenomenon was that clustered regularly interspaced short palindromic repeat (CRISPR) elements in bacteria conferred resistance in a cas-protein dependent manner.^3,10^ Since then, experimental results aiming to induce CRISPR-mediated resistance in clinical strains of *C. acnes* have suggested inconsistency with the notion of CRISPR as a predominant immunity mechanism, indicating other mechanisms are also at play.^11^

An alternative cause of bacteriophage resistance is the mechanism of superinfection exclusion (SIE). SIE is a property conferred by phage proteins, often from genes expressed by prophages, that gives rise to superinfection resistance (SIR).^12–14^ This is the case for phages like T7, T4, lambda, P1, and P2 which infect *Escherichia coli*, and phage P22 which infects *Salmonella typhimurium*.^15,16^ Phages like TP-J34, TP-778L, MG1363, and IL1403 have also been found to conduct SIE, targeting hosts including *Streptococcus thermophilis* and *Lactococcus lactis*.^13,17–19^ The SIE mechanism accounting for observed superinfection resistance among infected *C. acnes* bacteria has not been previously characterized. Therefore, it remains unknown whether there is a mechanism dependent on the expression of SIE gene(s) from a latent phage genome. *C. acnes* phages exhibit pseudolysogeny, an unstable state in which the phage genome remains as an episome and does not result in direct host cell lysis .^11,20^ This ability to maintain in the host cell supports the possibility of SIE as a phage resistance mechanism in *C. acnes*.

Previously, Liu *et al*. found that the bacteria that grew within the plaque centers exhibited SIR, as demonstrated by no lysis upon spotting these bacteria with the phage that was previously able to infect.^11^ In the present study, *C. acnes* phage Aquarius was isolated from the facial microcomedones of a donor without a history of acne. An early host range assay revealed a superinfection resistant phenotype in which bacterial growth was observed in the centers of plaque clearings on bacterial lawns of *C. acnes* ATCC 6919. This observation in phage Aquarius prompted a search for SIE characteristics, including verifying Aquarius’s ability to undergo pseudolysogeny, the only form of latency observed in *C. acnes* phages.^11,20^ Sequencing and genomic annotation of the Aquarius genome was performed and bioinformatics were used to characterize a putative SIE gene. Evidence suggesting a phage-mediated SIE phenotype could shed light on shortcomings of past phage-therapy experiments. It would also present a target parameter for future studies aimed at fine-tuning phage therapies for the purpose of controlling pathological *C. acnes* presence.

## Materials and Methods

### *C. acnes* culturing and Phage Isolation

*C. acnes* cultures were grown as described previously by Marinelli *et al*. by incubation at 37°C for three days under anaerobic conditions using the AnaeroPack System (Mitsubishi Gas Chemical Company, Tokyo, Japan).^3^ For all subsequent experiments, *C. acnes* plates were incubated under these conditions, unless otherwise noted. *C. acnes* bacteriophage isolates were isolated and purified from facial microcomedones using Bioré® pore strips (Kao USA Incorporated, Cincinnati, OH, USA) applied to the nose as described previously.^1,3^ Bacteriophages used in this study are described in **<u>Table 1</u>**.

**Table 1.** *C. acnes* bacteriophages used in this study.

| Phage Name | GenBank Accession No. | Bacterial Host | Experiments |
| --- | --- | --- | --- |
| Aquarius<sup>a</sup> | MF919491 | *Cutibacterium acnes* | All experiments |
| Lauchelly<sup>a</sup> | KR337650 | *Cutibacterium acnes* | Superinfection; bioinformatics |
| BruceLethal<sup>a</sup> | KR347352 | *Cutibacterium acnes* | Superinfection |
| QueenBey<sup>a</sup> | KR347355 | *Cutibacterium acnes* | Superinfection |
| ATCC 29399B_C<sup>b</sup> | JX262225 | *Cutibacterium acnes* | Superinfection; escape mutant isolation |
| P100A<sup>c</sup> | JX262221 | *Cutibacterium acnes* | Superinfection |
| P100D<sup>c</sup> | JX262220 | *Cutibacterium acnes* | Superinfection |
| P104A<sup>c</sup> | JX262218 | *Cutibacterium acnes* | Superinfection |
| P105<sup>c</sup> | JX262219 | *Cutibacterium acnes* | Superinfection |
| TP-J34<sup>d</sup> | HE861935 | *Streptococcus thermophilus*;<br>*Lactococcus lactis* | Bioinformatics |
| TP-778L<sup>e</sup> | HG380752 | *Streptococcus thermophilus*;<br>*Lactococcus lactis* | Bioinformatics |

<sup>a</sup> Source: UCLA Advanced Research in Virology Undergraduate Laboratory Curriculum.<sup>29</sup>
<sup>b</sup> Source: Clear/lytic (C) plaque isolated by Marinelli *et al.* (2012) from a mixed population of clear and turbid plaques observed from *C. acnes* phage stock ATCC 29399B originally described by Webster and Cummins (1978).<sup>3,28</sup>
<sup>c</sup> Source: Marinelli *et al.* (2012)<sup>3</sup>
<sup>d</sup> Source: Neve *et al.* (2003)<sup>30</sup>
<sup>e</sup> Source: Ali *et al.* (2014)<sup>19</sup>

### Viral DNA Purification and Sequencing

Phage DNA was treated with 5 mg/mL DNase1 and 10 mg/mL RNAseA for 30 minutes. Viral DNA was extracted using the Promega Wizard® DNA Clean-Up System (Madison, Wisconsin) as described by the manufacturer’s protocol. The concentration of the isolated phage DNA was obtained using a NanoVue spectrophotometer (GE Healthcare, Chicago, Illinois). The phage genomes were sequenced using an Illumina MiSeq (Illumina, San Diego, CA, USA) and assembled using the software program Newbler (Roche, Branford, Connecticut, USA) at the Pittsburgh Bacteriophage Institute in Pennsylvania as described previously.^21^

### Superinfection Resistance Test

Lawns of *C. acnes* ATCC 6919 and three clinical isolates (strains 060PA1, 110PA3, and 020PA1; described in Fitz-Gibbon *et al*.) were inoculated with phage lysates and observed for bacterial regrowth within the plaques.^1^ Putative lysogens were collected and inoculated in RCM and incubated for three days at 37°C under anaerobic conditions. The putative lysogens were plated on A-Media and 10-fold dilutions of phage lysates were spotted on the lawns to test for superinfection resistance.

### Lysogen Patch Test

Isolated *C. acnes* bacteria displaying SIR phenotypes were tested to determine if they were lysogens. *C. acnes* ATCC 6919 was plated on A-Media hard agar via the soft agar overlay technique and the putative lysogens were then streaked onto these plates and monitored for spontaneous phage release after incubation. Negative control plates were prepared for each putative lysogen by streaking the bacterial samples on plates without ATCC 6919 to ensure the growth of the streaked bacteria. For the stability patch tests, the same techniques were employed while serially streaking the individual strains onto a lawn of ATCC 6919 bacteria every three days over the course of approximately six months.

### Pseudolysogeny PCR

Pseudolysogeny PCR as described by Liu *et al*. was performed to identify if the isolated lysogens harbored the Aquarius genome.^11^ A master mix was prepared using the forward primer 5’-CCG AAG CCG ACC ACA TCA CAC C-3’ and the reverse primer 5′-TCA TCC AAC ACC TGC TGC TGC C-3’. DNA from uninfected bacteria and DNA-free negative controls were also assayed. All amplicons were run on a 0.8%-0.9% agarose gel for 25 minutes at 100 V.

### Ltp-like protein identification

BLASTp was conducted against the Phagesdb.org database using the sequence of Ltp from phages TP-J34 and TP-778L.^18,19,22,23^ The EMBL-EBI protein sequence and classification tool InterPro was then used to analyze the signature profiles of Ltp from phages TP-778L and TP-J34, as well as gp41 of phages Lauchelly and Aquarius.^24^ Multiple Em for Motif Elicitation (MEME) was employed to search for putative motifs in the non-cytoplasmic domains identified by InterPro and BLASTp.^25^ The protein alignment and phylogeny analysis tool Mega7 was used to identify charge conserved residues within the putative active site domains of the *C. acnes* Ltp-like proteins by conducting a protein alignment of Ltp_TP-778L_ and Ltp_TP-J34_ with several *C. acnes* phage gp41 proteins.^26^ The web portal for protein structure and function prediction RaptorX was used to compare overall predicted disorder between gp41 and Ltp.^27^

### Escape Mutant Isolation

Isolation of escape mutants was performed by incubating 10 μL of phage Aquarius lysate or phage ATCC 29399B_C (Genbank Accession JX262225) lysate with bacterial lysogen strains for 30 minutes, followed by plating for lawns using the soft agar overlay technique. Lysate dilutions of 10^−0^ and 10^−1^ were used as experimental groups, with dilution spots of 10^−2^, 10^−4^, 10^−6^, and 10^−7^ on a lawn of ATCC 6919 as a control. Phage ATCC 29399B_C isolated from a mixed phage population by Marinelli *et al*. from an original ATCC 29399B stock.^3,28^

### *C. acnes* Phage Infection and qPCR

To analyze RNA levels at various stages in the infection cycle, bacterial cultures of ATCC 6919 and an Aquarius pseudolysogen were grown in RCM and diluted to an OD_600_ of 0.2. Phage-free ATCC 6919, the pseudolysogen, and ATCC 6919 plus Aquarius at a multiplicity of infection of 10 were incubated at 37°C for the 90-minute duration of the active infection period. Total RNA was isolated using a Qiagen RNeasy® Kit (QIAGEN Group, Valencia, CA, USA) and analyzed for purity via Bioanalyzer. Following the isolation, cDNA was generated from the RNA according to the Bio-Rad iSCRIPT cDNA kit protocol using a Bio-Rad thermocycler (Bio-Rad Laboratories, Inc.; Hercules, CA, USA). After obtaining cDNA, qPCR was performed on 2 μL of undiluted cDNA using the Roche KAPA SYBR FAST qPCR kit using the primers in **<u>Table 2</u>**.

**Table 2.**
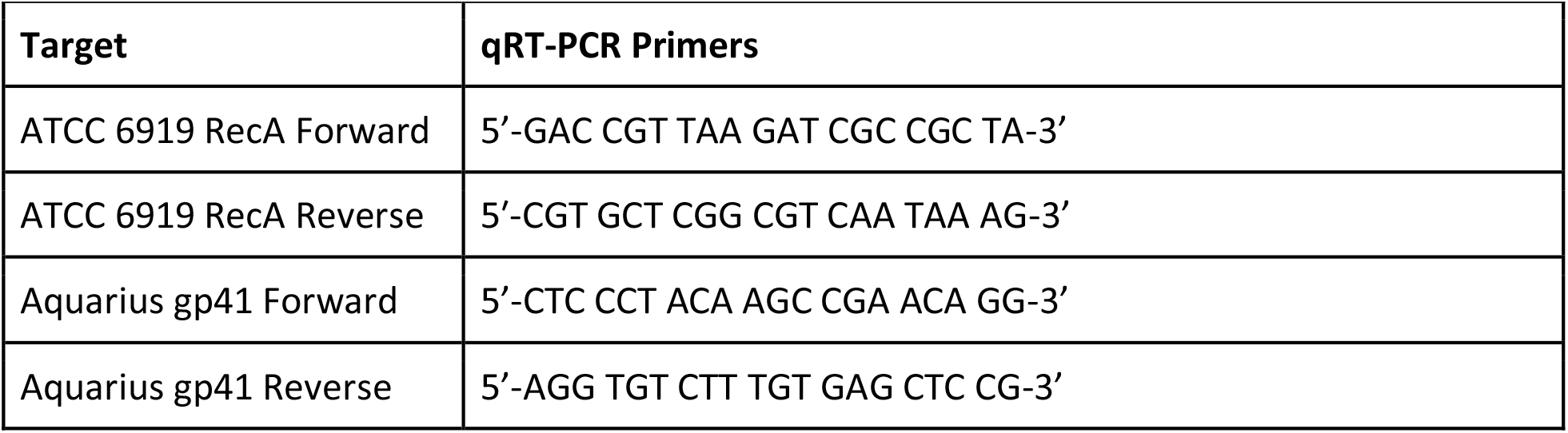
qRT-PCR primers for expression level analysis.

## Results and Discussion

### Phage Aquarius

The Aquarius genome was found to be 30112 bp with an 11 base 3’ sticky overhang (TCGTACGGCTT) and 54.5% GC content. Forty-eight putative ORFs were identified, twenty-nine of which were assigned a putative function. These genome characteristics were consistent with previously isolated *C. acnes* phages, and the 3’ sticky overhang indicates the genome is capable of the circularization necessary for pseudolysogeny.^3,11,20^

### Superinfection resistance

Bacterial growth was observed in the center of areas of phage Aquarius plaques on *C. acnes* ATCC 6919. To assay for superinfection resistance, bacteria were isolated from plaque centers, cultured, and used for spot tests with ten-fold dilutions of the Aquarius phage lysate. The spot tests revealed no evidence of lysis on the lawn of previously infected bacteria, suggesting SIR. Additionally, putative Aquarius lysogens of three *C. acnes* clinical isolates demonstrated superinfection resistance by phage Aquarius (**<u>Table 3</u>**). To assess the range of cross-immunity that the SIR mechanism may confer, eight additional representative *C. acnes* phages were spot tested for their ability to infect putative Aquarius lysogens; no lysis was observed on the lawns for any tested *C. acnes* phages (Table 2). To further test the range of this SIR mechanism, putative ATCC 6919 lysogens for seven different *C. acnes* phages were isolated and cross-tested for superinfection resistance for a variety of phages (**<u>Table 3</u>**). No lysis was observed for any of the tested combinations of putative lysogens and re-infecting phages, suggesting a mechanism for broadly preventing superinfection by *C. acnes* phages. *C. acnes* phages have been previously reported to have little genomic diversity, which may account for the effectiveness of the SIR mechanism.^3,11^

**Table 3.** Superinfection Immunity cross-infection summary.

| Infecting Phage | Bacterial Strain | Superinfection Test Phages |
| --- | --- | --- |
| Aquarius | ATCC 6919 | Aquarius, P100A, P100D, P104A, P105, Lauchelly, BruceLethal, QueenBey, ATCC 29399B_C |
| Aquarius | 060PA1, 110PA3, 020PA1<sup>1</sup> | Aquarius |
| P100A | ATCC 6919 | Aquarius, P100A, P100D, P104A, P105 |
| P100D | ATCC 6919 | Aquarius, P100A, P100D, P104A, P105 |
| P104A | ATCC 6919 | Aquarius, P100A, P100D, P104A, P105 |
| P105 | ATCC 6919 | Aquarius, P100A, P100D, P104A, P105 |
| Lauchelly | ATCC 6919 | Aquarius, Lauchelly, BruceLethal, QueenBey |
| BruceLethal | ATCC 6919 | Aquarius, Lauchelly, BruceLethal, QueenBey |
| QueenBey | ATCC 6919 | Aquarius, Lauchelly, BruceLethal, QueenBey |

<sup>1</sup> Source: All three clinical isolates described in Fitz-Gibbon *et al.* (2013)

### Pseudolysogeny

To test whether the SIR mechanism was associated with pseudolysogeny, putative Aquarius pseudolysogens were streaked onto an uninfected bacterial lawn. Areas of clearing were observed surrounding the streaks, indicating spontaneous phage release following lytic induction of the pseudolysogenized phage. PCR primers that anneal to the end of *C. acnes* phage genomes were used to assess if circularized phage genomes were present in the bacterial samples. Gel electrophoresis showed characteristic bands at roughly 735 base pairs for all putative pseudolysogens, and no bands on the uninfected ATCC 6919, consistent with previous experiments.^11^ This supports Aquarius’s capacity to undergo pseudolysogeny while simultaneously conferring SIR, as has been observed in *C. acnes* phages of previous studies.^20^

### Stability of pseudolysogens

Pseudolysogens are generally known to be less stable than full lysogens.^20^ Given this, the lytic centers (small non-plaque-like spots) scattered on pseudolysogen lawns may be a manifestation of induction of the lytic life cycle in bacteria after sequential passaging. Sub-isolates of two pseudolysogens identified in the previous experiment to have either a relatively high or low proportion of spontaneous lytic phage release, were assessed for pseudolysogen stability via patch testing for phage release over the course of several months. The sub-isolates of one lysogen, designated the “low stability” group, lost the ability to lyse the surrounding lawn after six total passages. The “high stability” group, however, sustained lytic capacity over six months of passaging.

Pseudolysogeny PCR was performed on both groups as described above to assay for maintenance of the circularized genome. All 27 high stability group sub-isolates produced a 735 bp band indicative of the presence of the phage genome, while the two representative samples for the low stability group did not, indicating phage genome loss. A Fisher’s exact test was conducted to assess independence between PCR result and patch test phenotype. The p-value was found to be *p* < 0.002, indicating a strong association between the two results.

Additionally, viral spot testing on cultures of the high and low stability groups after reaching five months of passaging revealed lysis of the low stability group, but sustained superinfection resistance by the high stability group. These results support the hypothesis of a pseudolysogeny associated phage resistance mechanism.

### Identification of Ltp-like protein

Bioinformatics was used to characterize putative gene(s) that may be involved in phage-mediated SIR. Previous research has shown that the Ltp protein in *Streptococcus thermophilus* and *Lactococcus lactis* phages TP-J34 and TP-778L confers SIR via an electrostatic interaction between Ltp and the tape measure protein C-terminus, resulting in a stalling of the ejection complex and prevention of infection.^18,19^ The Ltp_TP-J34_ and Ltp_TP-778L_ BLASTp yielded a hit with a gene of unknown function (gp41) in *C. acnes* phages Aquarius and Lauchelly, as well as to genes in *Propionibacterium freudenreichii* phages PFR1 and PFR2 (<u>Table 4</u>).^22^

**Table 4.** BLASTp results for Ltp-like proteins.

| Phage | Host | Ltp<sup>TP-J34</sup> Score | Ltp<sup>TP-J34</sup> E-Value | Ltp<sup>TP-778L</sup> Score | Ltp<sup>TP-778L</sup> E-Value |
| --- | --- | --- | --- | --- | --- |
| Aquarius | *C. acnes* | 26 | 9.7 | 31 | 0.4 |
| Lauchelly | *C. acnes* | 30 | 1.1 | 31 | 0.83 |
| PFR1 | *P. freudenreichii* | 110 | 6e-25 | 108 | 4e-24 |
| PFR2 | *P. freudenreichii* | 110 | 6e-25 | 108 | 4e-24 |

InterPro was used to compare signatures between Ltp_TP-J34_ and gp41 of Lauchelly and Aquarius.^24^ There was a remarkably similar signature profile between gp41 and Ltp, most noticeably the conservation of a long non-cytoplasmic domain at a similar locus and relative length within the sequences, as well as several signal peptide signatures (<u>Figure 1</u>). A roughly 90 amino acid-long region of disorder was also predicted in Ltp and gp41 by InterPro and RaptorX corresponding to the region after the end of the signal peptide at its C-terminal region and the beginning of the region within the non-cytoplasmic domain with which the first HTH domain of Ltp begins (<u>Figure 1</u>).^24,27^ MEME was employed to search for putative motifs within these non-cytoplasmic domains, considering a hallmark of Ltp family proteins is the presence of two repeat HTH domains (<u>Figure 1</u>).^18,25^ The output for Ltp_TP-J34_ and gp41 from sixteen *C. acnes* phages demonstrated two motifs of similar size with low p-values overlapping the non-cytoplasmic domain. Lastly, MEGA7 was used to identify charge conserved residues within the putative active site domains by conducting a protein alignment of Ltp_TP-778L_ and Ltp_TP-J34_ with several *C. acnes* phage gp41 protein products.^26^ The output indicated the presence of several charge conserved residues that may fit with the model of the Ltp-Tape Measure Protein (TMP) interaction according to Bebeacua *et al*., although further study is required to make definitive claims about active roles for any specific residue(s).^18^

**Figure 1.**
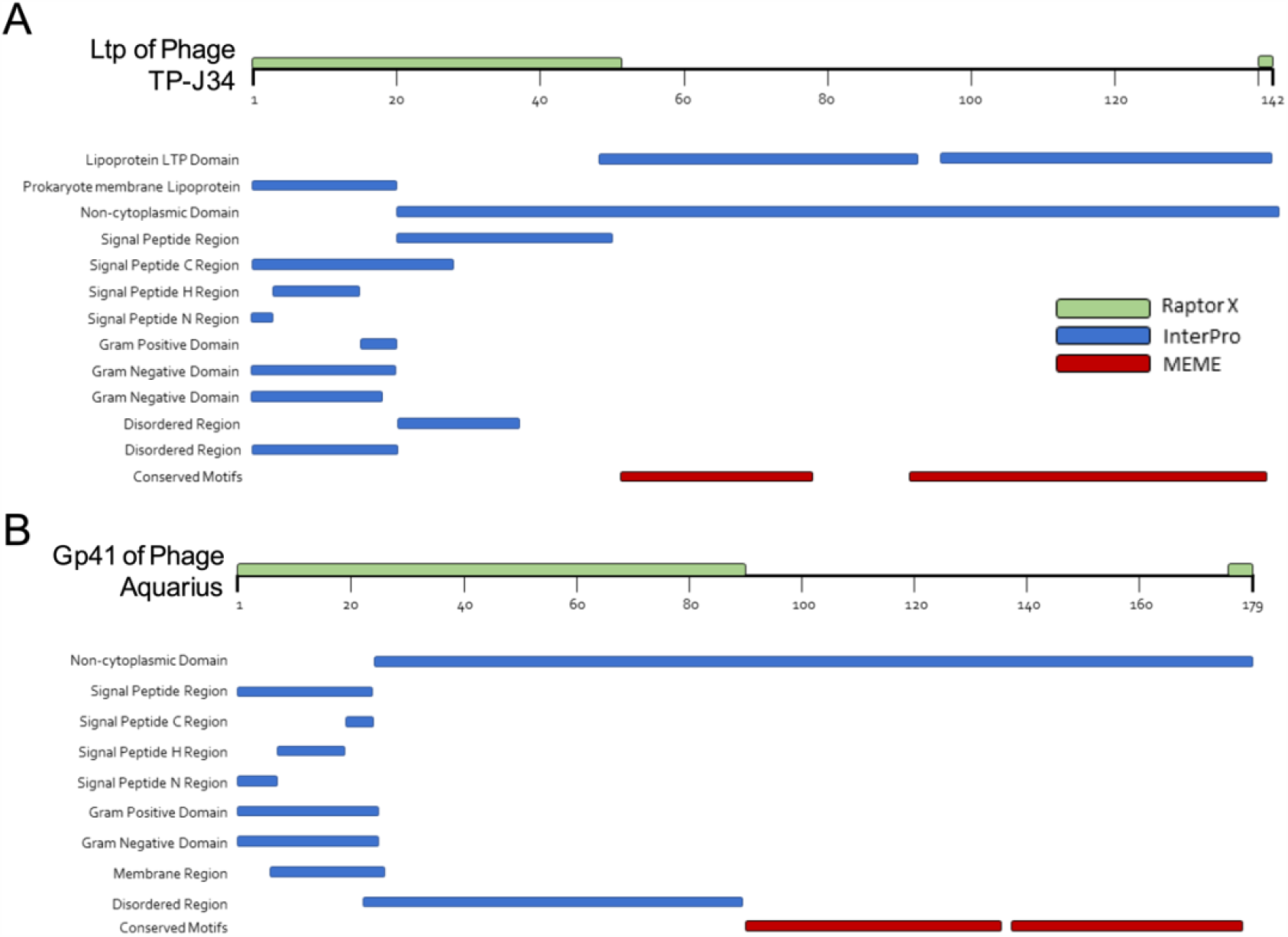
Protein signatures of Phage TP-J34 Ltp and Phage Aquarius Gp41. **(A)** *Ltp of Phage TP-J34*. Output from InterPro (blue) indicated the presence of several protein signatures, including the two conserved domains that comprise the active site region of Ltp (residues 49-92 and 96-141). Notable other signatures called included a prokaryotic lipoprotein (residues 1-20), regions of disorder (residues 21-50 and 21-37), signal peptide H-region (residues 4-15), signal peptide C-region (residues 16-20), signal peptide N-region (residues 1-3), signal peptide (residues 1-20), transmembrane signal peptide (residues 1-28), and a non-cytoplasmic domain (residues 21-142). Output from MEME (red) indicated the presence of two conserved motifs as well, spanning residues 46-77 and 91-141. Output from RaptorX (green; depicted on ruler) also identified a generally high region of disorder spanning from the first residue to roughly residue 50, and a small region at the very end of the peptide spanning roughly one to two residues. **(B)** Gp41 of Phage Aquarius. Output from InterPro (blue) indicated the presence of several notable protein signatures, including regions of disorder (residues 23-89 and 34-51), signal peptide H-region (residues 8-19), signal peptide C-region (residues 20-24), signal peptide N-region (residues 1-7), signal peptide (residues 1-24), transmembrane signal peptide (residues 1-25), transmembrane helix (residues 7-26), and a non-cytoplasmic domain (residues 25-179). Output from MEME (red) indicated the presence of two conserved motifs as well, spanning residues 84-133 and 137-177. Output from RaptorX (green; depicted on ruler) also identified a generally high region of disorder spanning from the first residue to roughly residue 91, and a small region at the very end of the peptide spanning roughly three to five residues.

### No escape mutants found

Since bioinformatics suggested the presence of a conserved Ltp-like protein in Aquarius’s genome, an attempt was made to isolate escape mutant phages capable of infecting pseudolysogens. It was hypothesized that phage may acquire SIR-surpassing mutations that map to the tape measure protein of Aquarius, which would support a mechanism characteristic of that described by Bebeacua *et al*.^18^ However, all attempts at plating Aquarius on pseudolysogens were unsuccessful in producing plaques. The lack of plaques following multiple attempts supports the notion of a tight immunity mechanism at play.

### qPCR of Aquarius Ltp-like gp41

To find evidence linking gp41 to SIR, qPCR was performed to assess gene expression in actively infected bacteria compared to a phage-free control, and Aquarius pseudolysogens. The average results of three qPCR trials indicated a fold increase of approximately 333,000 times more expression of Aquarius gp41 in the active infection group and 40,000 times more expression in the pseudolysogen group relative to the control after normalizing samples with the bacterial housekeeping gene RecA. One-way ANOVA with post hoc Tukey HSD comparison of the qPCR results yielded a p-value of <0.05, suggesting an important role for gp41 early in phage infection. This suggests that gp41 is highly expressed during initial infection, at the period in which many individual phages have recently entered into corresponding hosts and making life cycle decisions, and expression is maintained at a lower level during the pseudolysogenic life cycle.

## Conclusion

The results of this research strongly support the previous findings that Aquarius and other *C. acnes* phages are able to undergo the pseudolysogeny, and supports the hypothesis of a superinfection resistance mechanism which is actively-transcribed from semi-stable episomes within their host. Bioinformatics demonstrates the presence of a protein in all *C. acnes* phage genomes tested which remarkably resembles the known SIR protein Ltp. At this time, it remains unclear whether the mechanism of gp41 in Aquarius is governed by a mechanism paralleling that of Ltp, which is characterized by negatively charged residues on the Ltp surface interacting with predominantly positively charged surface-peptides on TMP during phage DNA ejection into its host.^18^ Despite this, there is clear involvement of gp41 in the phage life cycle, given the magnitude of mRNA expression during active- and post-infection, as determined by qPCR. The lower gp41 expression in latent pseudolysogen samples relative to the sharp increase of the active infection samples may suggest the possibility of breaching a saturation-point or feedback mechanism, resulting in waning gp41 expression over time.

Lastly, given current constraints on cloning via electroporation in C. acnes, phage recombineering involving a gp41 knockout and complementation presents a future route for solidifying gene characterization. Should this demonstrate the association of superinfection resistance with gp41, x-ray crystallography may be warranted to solidify understanding of the gene’s mechanism in C. acnes phages on a bio-molecular level. Gp41’s mechanism should then be compared to the mechanisms of SIR in other phages to assess evolutionary similarities and novelty. An understanding of the mechanism(s) by which this resistance may be produced in bacteria that were previously capable of being lysed by phage may give rise to the exploitation of the phenomenon in the fine tuning of phage-based therapeutics. Likewise, this approach would give rise to a more thorough understanding of infection in the context of the evolutionary life cycles that the phage is capable of undergoing to promote its propagation and the success of its survival.

## Acknowledgements

The authors acknowledge the research support for the present publication that include: the SEA-PHAGES program at the University of California, Los Angeles (UCLA), the Microbiology, Immunology, and Molecular Genetics department at UCLA, the UCLA Dermatology department, and the UCLA Dean of the Life Sciences Division. The authors acknowledge the research support of Daniel Russel from the University of Pittsburg for his contributions regarding genome assembly and quality control. The authors also acknowledge the assistance of Dr. Kris Reddi from UCLA, as well as the intellectual assistance of Rocky Ng, Andrew Lund, Kindra Kelly-Scumpia, and Dr. Francie Mercer for their contributions to the study and assistance with quality control.

## Authorship Statement

SW conceived the project idea. JMP and LJM in conjunction with the SEA-PHAGES program conceived and designed the experiments. SW, SM, CS, and CN isolated phage Aquarius and performed phage characterization experiments. SW performed the superinfection and pseudolysogeny-related experiment and analyzed the data with the assistance of LJM. RM, LJM, and JMP contributed reagents and materials, as well as professional guidance. SW drafted the manuscript; SM, CS, and CN assisted with preparation of the materials and methods section of the paper, as well as edits and feedback. JPM edited the final manuscript. All co-authors have reviewed and approved of the manuscript prior to submission. This statement certifies that this manuscript has been submitted solely to this journal and is not published, in press, or submitted elsewhere.

## References

1. Fitz-Gibbon, S. et al. Propionibacterium acnes Strain Populations in the Human Skin Microbiome Associated with Acne. J Invest Dermatol 133, 2152–2160 (2013).

2. Li, H. The Human Skin Microbiome in Health and Skin Diseases. in Metagenomics of the Human Body (ed. Nelson, K. E.) 145–163 (Springer, 2011). doi:10.1007/978-1-4419-7089-3_8.

3. Marinelli, L. J. et al. Propionibacterium acnes Bacteriophages Display Limited Genetic Diversity and Broad Killing Activity against Bacterial Skin Isolates. mBio 3, e00279–12 (2012).

4. Perry, A. & Lambert, P. Propionibacterium acnes: infection beyond the skin. Expert Review of Anti-infective Therapy 9, 1149–1156 (2011).

5. Leheste, J. R. et al. P. acnes-Driven Disease Pathology: Current Knowledge and Future Directions. Front. Cell. Infect. Microbiol. 7, (2017).

6. Brüggemann, H. & Lood, R. Bacteriophages infecting Propionibacterium acnes. Biomed Res Int 2013, 705741 (2013).

7. Jończyk-Matysiak, E. et al. Prospects of Phage Application in the Treatment of Acne Caused by Propionibacterium acnes. Front. Microbiol. 8, (2017).

8. Coenye, T., Peeters, E. & Nelis, H. J. Biofilm formation by Propionibacterium acnes is associated with increased resistance to antimicrobial agents and increased production of putative virulence factors. Research in Microbiology 158, 386–392 (2007).

9. Holmberg, A. et al. Biofilm formation by Propionibacterium acnes is a characteristic of invasive isolates. Clinical Microbiology and Infection 15, 787–795 (2009).

10. Brüggemann, H., Lomholt, H. B. & Kilian, M. The flexible gene pool of Propionibacterium acnes. Mobile Genetic Elements 2, 145–148 (2012).

11. Liu, J. et al. The diversity and host interactions of Propionibacterium acnes bacteriophages on human skin. ISME J (2015) doi:10.1038/ismej.2015.47.

12. Labrie, S. J., Samson, J. E. & Moineau, S. Bacteriophage resistance mechanisms. Nature Reviews Microbiology 8, 317–327 (2010).

13. Sun, X., Göhler, A., Heller, K. J. & Neve, H. The ltp gene of temperate Streptococcus thermophilus phage TP-J34 confers superinfection exclusion to Streptococcus thermophilus and Lactococcus lactis. Virology 350, 146–157 (2006).

14. Seed, K. D. Battling Phages: How Bacteria Defend against Viral Attack. PLOS Pathogens 11, e1004847 (2015).

15. McAllister, W. T. & Barrett, C. L. Superinfection exclusion by bacteriophage T7. Journal of Virology 24, 709–711 (1977).

16. Hofer, B., Ruge, M. & Dreiseikelmann, B. The superinfection exclusion gene (sieA) of bacteriophage P22: identification and overexpression of the gene and localization of the gene product. Journal of bacteriology 177, 3080–3086 (1995).

17. Mahony, J., McGrath, S., Fitzgerald, G. F. & Sinderen, D. van. Identification and Characterization of Lactococcal-Prophage-Carried Superinfection Exclusion Genes. Appl. Environ. Microbiol. 74, 6206–6215 (2008).

18. Bebeacua, C. et al. X-ray structure of a superinfection exclusion lipoprotein from phage TP-J34 and identification of the tape measure protein as its target. Molecular Microbiology 89, 152–165 (2013).

19. Ali, Y. et al. Temperate Streptococcus thermophilus phages expressing superinfection exclusion proteins of the Ltp type. Front. Microbiol. 5, (2014).

20. Lood, R. & Collin, M. Characterization and genome sequencing of two Propionibacterium acnes phages displaying pseudolysogeny. BMC Genomics 12, 198 (2011).

21. Russell, D. A. Sequencing, Assembling, and Finishing Complete Bacteriophage Genomes. in Bacteriophages: Methods and Protocols, Volume 3 (eds. Clokie, M. R. J., Kropinski, A.M. & Lavigne, R.) 109–125 (Springer New York, 2018). doi:10.1007/978-1-4939-7343-9_9.

22. Altschul, S. F., Gish, W., Miller, W., Myers, E. W. & Lipman, D. J. Basic local alignment search tool. Journal of Molecular Biology 215, 403–410 (1990).

23. Russell, D. A. & Hatfull, G. F. PhagesDB: the actinobacteriophage database. Bioinformatics 33, 784–786 (2017).

24. Mitchell, A. et al. The InterPro protein families database: the classification resource after 15 years. Nucleic Acids Res 43, D213–D221 (2015).

25. Bailey, T. L. & Elkan, C. Fitting a mixture model by expectation maximization to discover motifs in biopolymers. Proc Int Conf Intell Syst Mol Biol 2, 28–36 (1994).

26. Kumar, S., Stecher, G. & Tamura, K. MEGA7: Molecular Evolutionary Genetics Analysis Version 7.0 for Bigger Datasets. Mol Biol Evol 33, 1870–1874 (2016).

27. Källberg, M. et al. Template-based protein structure modeling using the RaptorX web server. Nature Protocols 7, 1511–1522 (2012).

28. Webster, G. F. & Cummins, C. S. Use of bacteriophage typing to distinguish Propionibacterium acne types I and II. Journal of Clinical Microbiology 7, 84–90 (1978).

29. Shapiro, C. et al. Comparing the Impact of Course-Based and Apprentice-Based Research Experiences in a Life Science Laboratory Curriculum. J Microbiol Biol Educ 16, 186–197 (2015).

30. Neve, H., Freudenberg, W., Diestel-Feddersen, F., Ehlert, R. & Heller, K. J. Biology of the temperate Streptococcus thermophilus bacteriophage TP-J34 and physical characterization of the phage genome. Virology 315, 184–194 (2003).

